# Leaf trichome distribution pattern in *Arabidopsis* reveals gene expression variation associated with environmental adaptation

**DOI:** 10.1101/2020.07.01.180794

**Authors:** Shotaro Okamoto, Kohei Negishi, Yuko Toyama, Takeo Ushijima, Kengo Morohashi

## Abstract

Gene expression varies stochastically even in both heterogenous and homogeneous cell populations. This variation is not simply useless noise; rather, it is important for many biological processes. Unicellular organisms or cultured cell lines are useful for analyzing the variation in gene expression between cells; however, owing to technical challenges, the biological relevance of this variation in multicellular organisms such as higher plants remains unclear. Here, we addressed the biological relevance of this variation between cells by examining the genetic basis of trichome distribution patterns in *Arabidopsis thaliana*. The distribution pattern of a trichome on a leaf is stochastic and can be mathematically represented using Turing’s reaction-diffusion (RD) model. We analyzed simulations based on the RD model and found that the variability in the trichome distribution pattern increased with the increase in stochastic variation in a particular gene expression. Moreover, differences in heat-dependent variability of the trichome distribution pattern between the accessions showed a strong correlation with environmental factors to which each accession was adapted. Taken together, we successfully visualized variations in gene expression by quantifying the variability in the *Arabidopsis* trichome distribution pattern. Thus, our data provide evidence for the biological importance of variations in gene expression for environmental adaptation.

## 1. Introduction

A genetically identical population can exhibit phenotypic variation arising from subtle differences in gene expression [1,2]. Gene expression is modulated by intrinsic developmental cues and environmental stimuli, and stochastic fluctuations in gene expression are important for many biological processes including environmental stress response, survival, and adaptation [2,3]. Unicellular organisms (e.g., bacteria) or single cells (e.g., cultured mammalian cell lines) are useful for studying variation in gene expression between cells [4]. For instance, embryonic stem cells exhibit significant heterogeneity in gene expression [5].

Adaptation to specific environmental factors such as light and temperature is essential for plants and requires a system for controlling variation in gene expression. Recent studies have investigated stochastic variations in plant phenotype such as phyllotactic patterning and the timing of epidermal cell division in the sepal [6,7]. The findings of these studies suggest that such variations can be beneficial for plants, although the precise underlying mechanisms remain unknown [6,8–11].

Single-cell transcriptomics is currently the main method used to measure variation in gene expression between cells. However, single-cell transcriptomics is costly, and separating plant cells from organs requires enzymatic treatment, which can artificially alter gene expression profiles. Although variation in gene expression in higher order multicellular organisms was recently measured, differences of gene expression variation between genes has not yet been demonstrated [11]. The trichome, a hair-shaped organ that develops from a pluripotent epidermal cell at an early stage of leaf development, performs multiple functions including herbivore defense, leaf moisture retention, and metabolite secretion. Trichomes serve as a useful trait for studying environmental adaptation in plants. The distribution pattern of leaf trichomes in *Arabidopsis thaliana* is a suitable system for investigating variation in gene expression in plants for the following reasons. First, the trichome position on a leaf is equally and probabilistically distributed [10,12,13], thus, the trichome distribution pattern is believed to emerge stochastically. Second, the gene regulatory network related to trichome development has been well studied [14–16]; *GLABRA 3* (*GL3*), which encodes a basic helix-loop-helix transcription factor, is critical for cell fate determination during trichome formation. The GL3 protein forms a complex with GL1, an R2R3-MYB transcription factor, to control the expression of downstream genes. The GL3-GL1 complex activates not only positive regulators of trichome cell fate determination, but also R3-MYB transcription factors that directly interact with GL3 in neighboring cells to inhibit the formation of the GL3-GL1 complex. Consequently, these cells do not form trichomes. Third, the trichome distribution pattern is explained by Turing’s reaction-diffusion (RD) model [13,17,18], and a trichome distribution pattern was mathematically and experimentally demonstrated [19] (Fig. 1A). Since the pattern formation required stochastic fluctuations, the previously proposed RD model uniformly added a stochastic noise. However, in the model, the variations in individual factors were neglected. We speculated that differences in gene expression between epidermal cells immediately before trichome cell fate determination can affect the trichome distribution pattern (Fig 1B). To predict the effect of variations in gene expression between epidermal cells on trichome development, a proper mathematical model and further experiments are needed.

**Figure 1.**
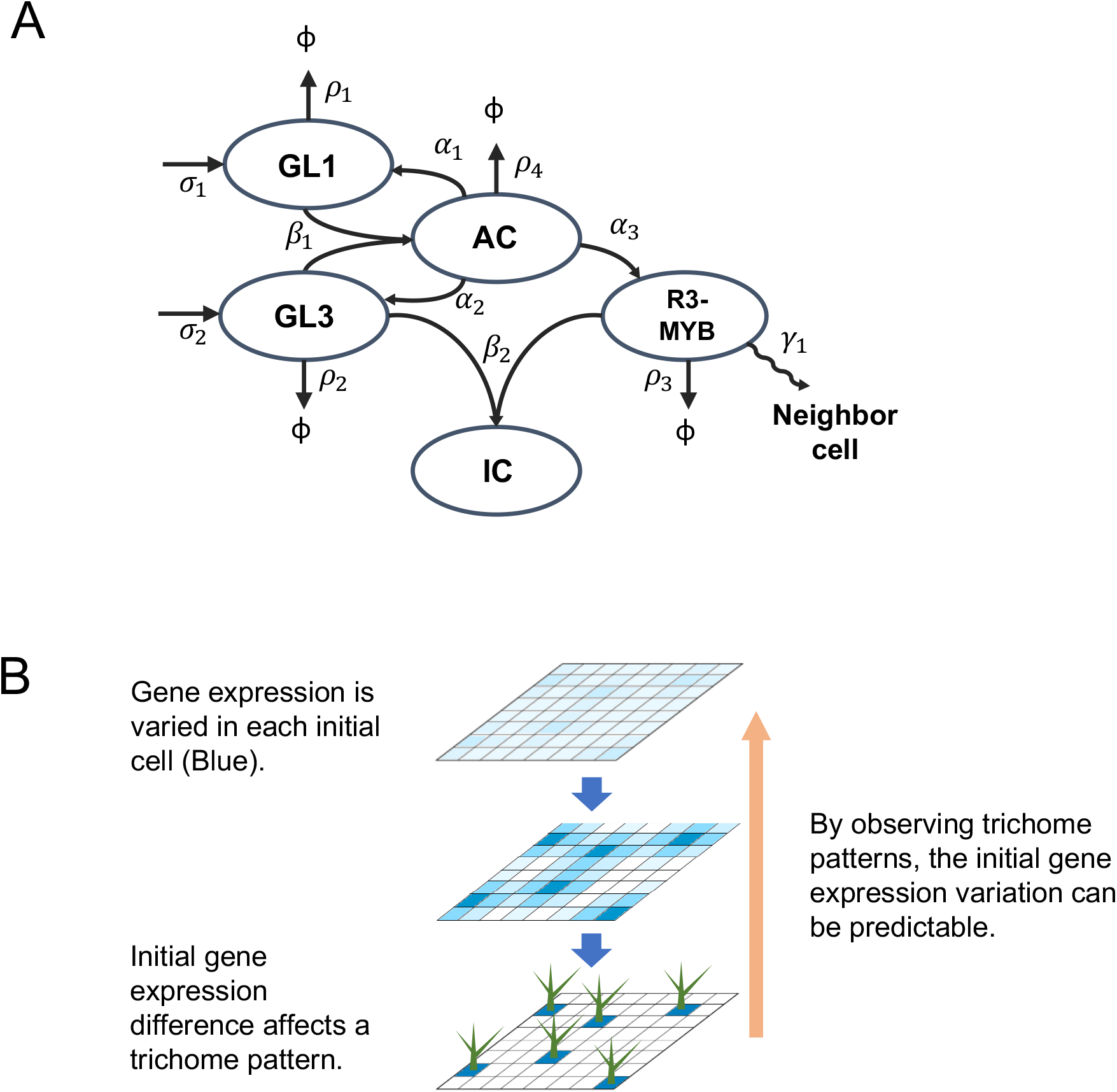
Concept of the hypothesis proposed in the text. A) Interaction scheme underlying trichome formation. GL1 and GL3 are expressed at rates of σ_1_ and σ_2_, respectively. GL1 and GL3 form an active complex (AC) at rate β1 to activate GL1, GL3, and R3-MYB at rates α1, α2, and α3, respectively. GL3 and R3-MYB interact at rate β2 to form an inactive complex (IC). GL1, GL3, AC, and R3-MYB are degraded at rates ρ1, ρ2, ρ4, and ρ3, respectively. The R3-MYB complex moves to neighboring cells at rate γ1. B) Schematic representation of the hypothesis proposed in the text. Gene expression variation between cells is represented by the intensity of the blue color (top). Gene expression variations increased (middle), and trichomes eventually formed based on the initial gene expression variations (bottom). According to our hypothesis (orange), gene expression variations in each initial cell can be predicted by measuring trichome distribution patterns.

In this study we investigated the factors affecting the range of variability in gene expression between individual cells, and by performing experiments using fresh leaf material. We visualized heterogeneity in GL3 expression, which correlated with the regularity of the trichome distribution pattern. We found that variations in gene expression were affected by histone modifications and ambient temperature, and were correlated with the mean annual temperature of the habitat of *A. thaliana* accessions. While our approach consists of both intrinsic and extrinsic variations, our model enables the quantification of gene expression variation and provides a basis for investigating the role of gene expression variation in environmental adaptation.

## 2. Results

### 2.1 Computational simulation of the effect of variation in gene expression on trichome patterning

We hypothesized that the leaf trichome distribution pattern is determined by differences in gene expression between individual epidermal cells. To test this hypothesis, we first established a method for quantifying the trichome distribution pattern. Trichome positions on a mature leaf are stable; i.e., the distribution of trichomes on a leaf is considered as a regular pattern. We measured the distance between the two closest trichomes (next-neighbor distance [ND]) and examined the distribution of NDs of all trichomes on a leaf. A truly regular trichome pattern shows a single ND distribution peak since the shape formed by connecting the positions of three trichomes is an equilateral triangle. On the other hand, the distribution of NDs broadens as the trichome pattern becomes more irregular. We calculated the variance in the distribution of NDs and carried out a quantitative comparison between leaves based on the normalized ND (NND) value, calculated by dividing ND with the average distance between trichomes on a leaf (Fig 2).

**Figure 2.**
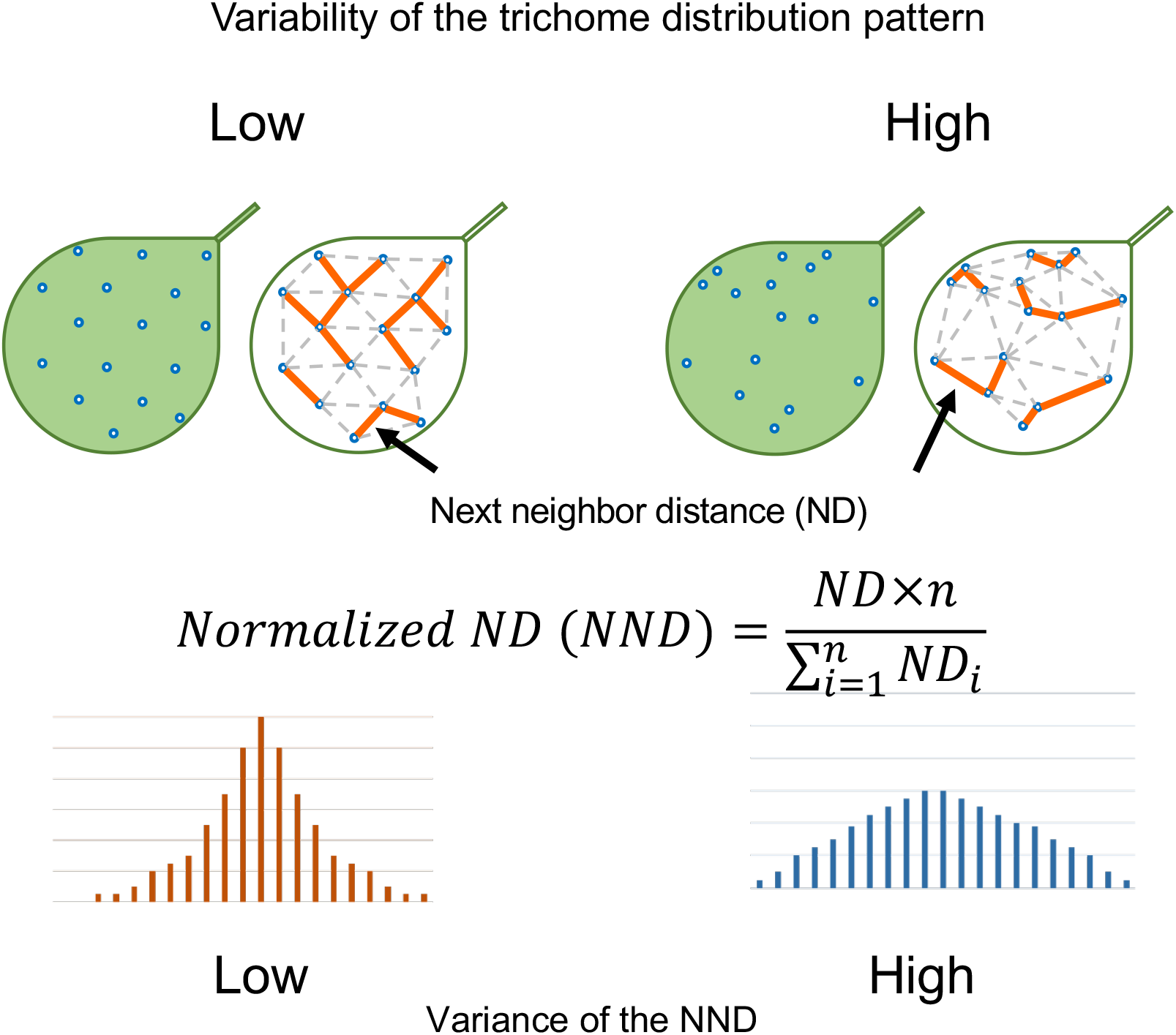
Theoretical interpretation of the quantification of the trichome distribution pattern. The orange line indicates the shortest path between trichomes, referred to as next-neighbor distance (ND). The equation used to calculate the normalized ND (NND) is shown in the middle. The histogram of NNDs is shown at the bottom. High variability of the trichome distribution pattern demonstrates high variance of the NND distribution

We applied the NND quantification method to computational simulations before performing experiments using real leaves. In the previous RD model [13], differential equations were solved using fixed parameters, followed by application of a 1% variation to all cells. However, this calculation did not apply a specific variation to each parameter of a single gene, which is unlikely to reasonably reflect a biological system. Therefore, we separately applied a variation to each parameter. That is, we applied two different variations: equal variations to all cells as in the previous model [13] and specific variations applied separately to individual parameters. In the previous model, there were 14 dimensionless parameters. To facilitate interpretation, we used parameters before the dimensionless treatment (Table 1). We anticipated that our modified mathematical model would distinguish gene-specific variation from natural stochastic variation. We applied a maximum variation of 50% of the values of parameters to our modified model. To evaluate whether 0–50% variation was a suitable range, we analyzed publicly available single-cell RNA sequencing data [20]. The coefficient of variation (CV) of gene expression between 13 single cells was 1.12 (red line in Fig. S1). In our simulations, a range of CVs by applying 0–50% variations was between 0.05 and 0.2, which was far below 1.12. Although Fig S1 shows CVs of only GL3, all CVs from other components were less than 1.12. These data indicate that the applied range is biologically relevant. We also found that the number of trichomes and the regularity of their patterns were not correlated (Fig. S2).

**Table 1.** Parameters used in this study.

| Parameter | Function | Value |
| --- | --- | --- |
| $\sigma_1$ | Basal GL1 transcription rate | 8.2707 |
| $\alpha_1$ | AC-regulated GL1 transcription rate | 3.4869 |
| $\rho_1$ | GL1 degradation rate | 1 |
| $\beta_1$ | GL1-GL3 interaction rate | 1 |
| $\sigma_2$ | Basal GL3 transcription rate | 15.0952 |
| $\alpha_2$ | AC-regulated GL3 transcription rate | 1.3488 |
| $\rho_2$ | GL3 degradation rate | 0.4503 |
| $\beta_2$ | GL3-R3MYB interaction rate | 7.9509 |
| $\alpha_3$ | AC-regulated R3MYB transcription rate | 0.4117 |
| $\rho_3$ | R3MYB degradation rate | 0.9565 |
| $\gamma_1$ | R3MYB transport rate | 9.565 |
| $\rho_4$ | AC degradation rate | 0.2703 |

In this study, the F test was used to assess the statistical significance of differences in variance as compared with no particular variation in each parameter. After 500 trials, our simulations showed that both the regularity of trichome patterns and the number of trichomes were affected in some parameters (Fig. S3). Since we focused on the parameters that influenced only the regularity of the trichome distribution pattern, we considered the parameters that affected only variances and not the number of trichomes, and ignored other parameters that affected variances of NND distributions. There are three types of parameters in the model; σ, which represents the rate of gene expression; ρ, which represents the rate of degradation; and β, which represents the rate of protein-protein association. Consequently, σ_2_ was the only parameter that increased the variance of NNDs as the variation of the parameters increased, suggesting that a change in σ_2_ (rate of *GL3* expression) affects the regularity of the trichome distribution pattern (Fig. 3). It is worth noting that σ_1_ (rate of *GL1* expression) did not affect this pattern (Fig. 3), even though GL1 is a key component of the active GL1-GL3 complex that controls downstream genes modulating trichome cell fate determination [14,16]. These results suggest that the regularity of the trichome distribution pattern reflects variations in GL3 expression according to the mathematical model.

**Figure 3.**
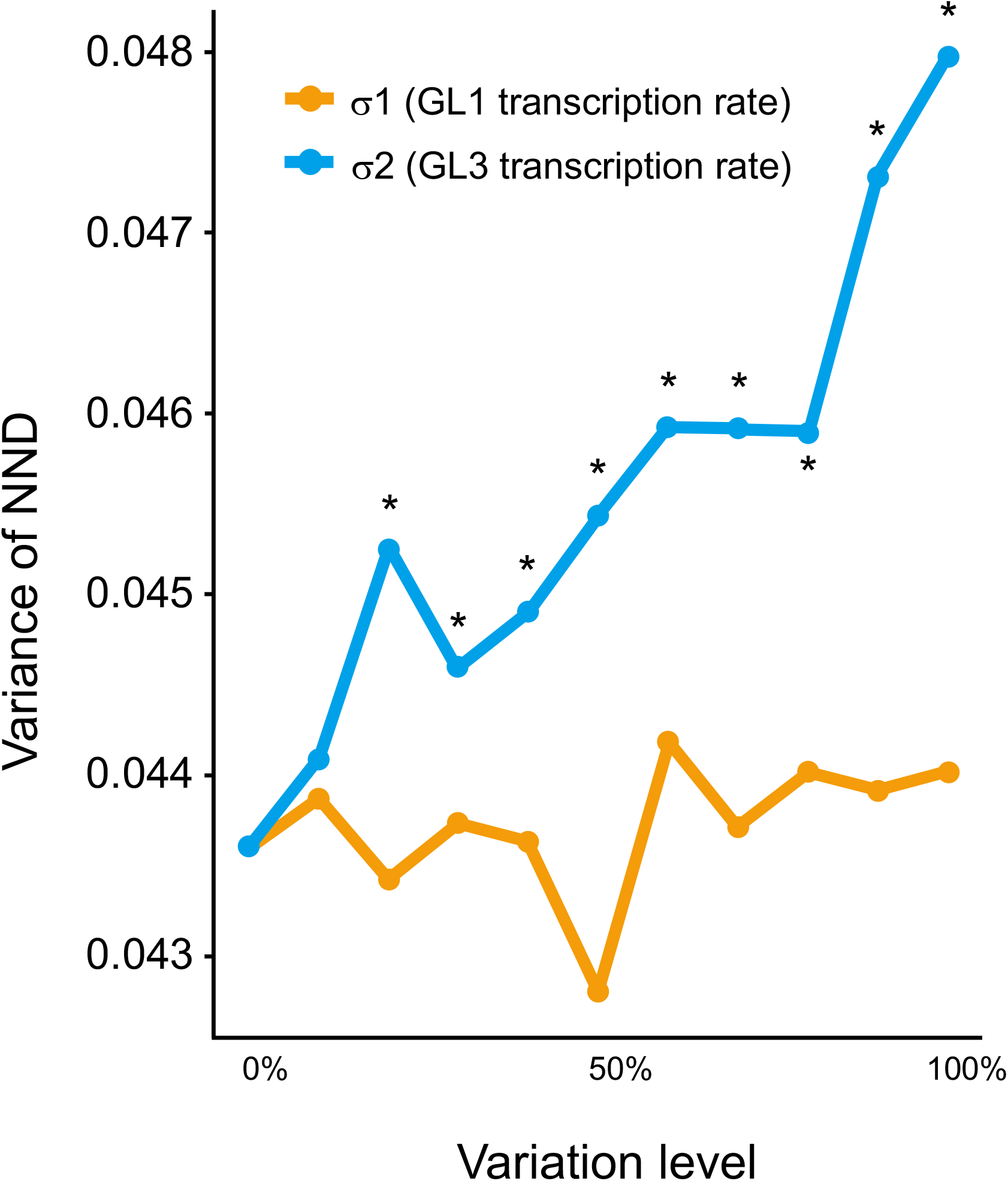
Effects of the degree of variation in each parameter. Variances of normalized next-neighbor distance (NND) increased proportionally with variations in σ2, whereas σ1 was unaffected.

### 2.2 The trichome distribution pattern is independent on the formation of other epidermal cell patterns

To validate our mathematical model, we analyzed trichome patterns on the third and fourth mature leaves collected from 3-week-old *A. thaliana* plants. NNDs were quantified using a method described previously [21]. Briefly, each sampled leaf was cleared by incubating in 80% lactic acid. Since a trichome cell walls exhibit polarizing (birefringent) properties, we used polarized light microscopy (PLM) to distinguish trichomes from non-trichome epidermal cells, followed by automated analysis to determine the trichome distribution patterns of all sampled leaves.

Since trichomes develop from an epidermal cell, their distribution pattern is associated with the pattern of non-trichome cells such as pavement cells and stomata cells on a leaf. The distribution of stomata is more deterministic than that of trichomes and is unlikely to follow Turing’s RD model [22]. It was previously reported that the Voronoi diagram based on stomata position is correlated with the shape of a pavement cell, which is accounted for by mechanochemical polarization of contiguous pavement cell walls [23,24]; we therefore assumed that the stomata pattern reflects the pattern of pavement cell shapes and that patterns of trichomes and other epidermal cells can be distinguished by simultaneously measuring those of trichome and stomata positions.

To demonstrate that the stomata pattern is independent of the trichome distribution pattern, we analyzed the pattern of pavement cells by fluorescence microscopy following propidium iodide (PI) staining. We also examined epidermal cell patterns in the *repressor of lrx 1*(*rol1*) mutant [25], in which cell wall composition is perturbed, resulting in rounder pavement cells compared with the wild type. We measured both trichome and stomata patterns of leaves harboring two alleles of the *rol1* mutant (*rol1-1* and *rol1-2*). No correlation was observed between the NND variances of stomata and trichomes (Fig. 4). The variances of stomata NNDs were reduced in *rol1* mutants, along with a corresponding reduction of in the variability of pavement cell patterns; however, no difference between the wild type of *rol1-1* mutant was observed in trichome patterns. These results suggest that trichome and pavement cell patterns are established independently of each other.

**Figure 4.**
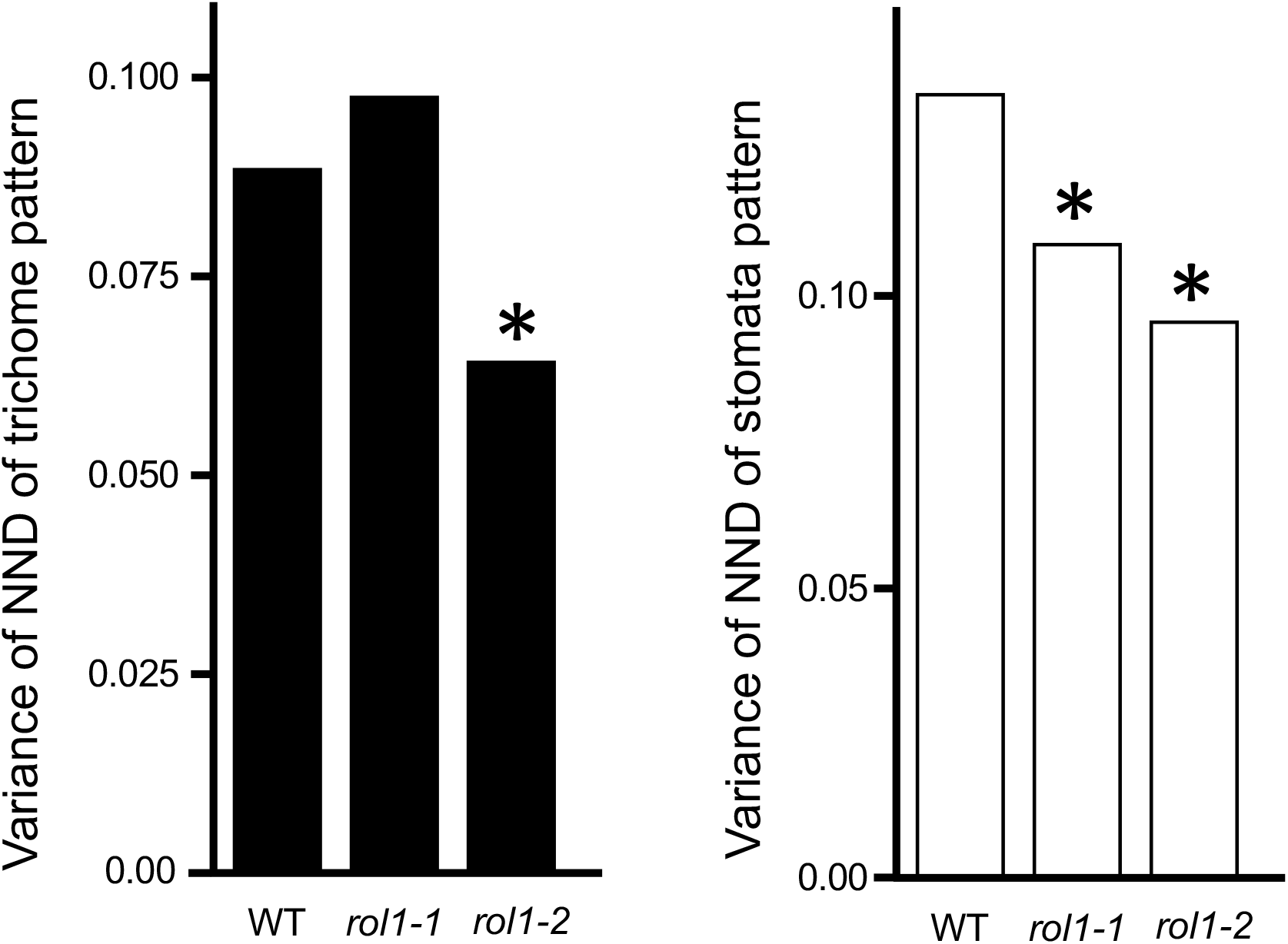
Normalized next-neighbor distance (NND) variances of trichomes and stomata of Columbia (Col) and rol1 mutant. Two alleles of the rol1 mutant showed increased regularity of stomata patterns (white); however, the trichome pattern showed no correlation (black). *P < 0.05 (F test).

### 2.3 Gene expression variations between cells increase at elevated temperatures

Given that environmental stimuli such as light and temperature, can alter gene expression [26], we investigated whether modest changes in light intensity or temperature would affect gene expression variations. Exposure of seedlings to a relatively strong light intensity perturbed both trichome and stomata distribution patterns (Fig. S4). However, seedlings exposed to a modestly high temperature (26°C) showed perturbed trichome distribution patterns but normal stomata distribution patterns compared with seedlings grown in plates at 22°C (Fig. 5), suggesting that gene expression was more varied at 26°C. A further increase in temperature to 30°C eventually disturbed stomata distribution patterns. These results suggest that the trichome distribution pattern is more sensitive to environmental fluctuations than the stomata distribution pattern.

**Figure 5.**
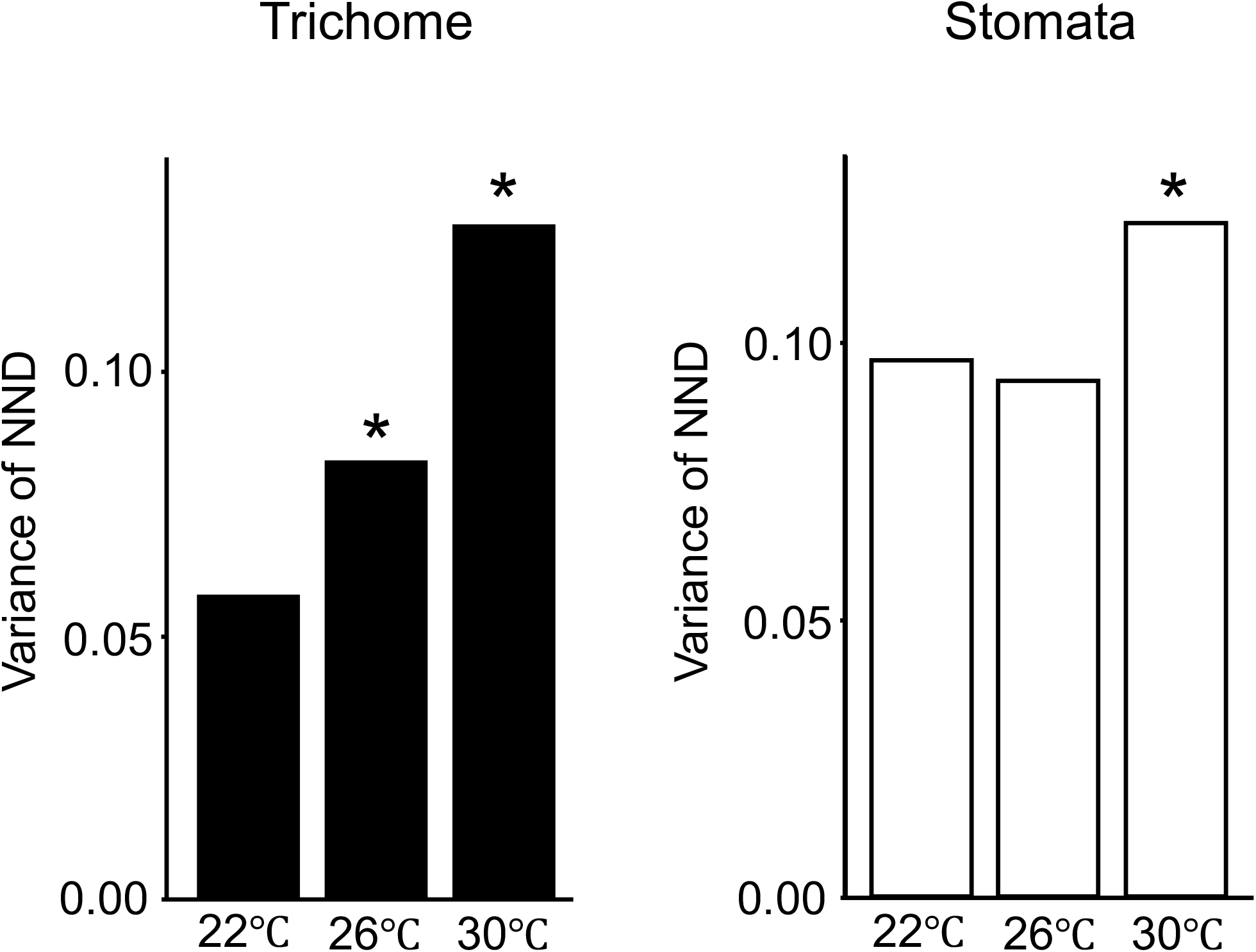
Normalized next-neighbor distance (NND) variances of trichomes and stomata of Col plants grown under mild heat stress. Variances of trichomes (black) and stomata (white) of Col plants grown at 22°C, 26°C, and 30°C are shown. NND variances of trichomes increased with the increase in temperature (black). *P < 0.05 (F test)

The results obtained from sampled leaves are likely to reflect the amounts of *GL3* gene products. To confirm that the variation of the GL3 protein amounts correspond to the NND distribution pattern, we evaluated the GL3 protein level in transgenic plants expressing the *GL3* gene fused to *yellow fluorescent protein* (*YFP*) gene under the control of the *GL3* promoter (GL3-YFP) [14,27]. Since we considered the trichome pattern but not mature trichomes per se, we measured YFP signals in the cells within the trichome initiation zone [28]. Variability in the intensity of GL3-YFP fluorescence in *Arabidopsis* leaves increased at 26°C (Fig. 6). The distribution of GL3-YFP fluorescence showed no significant difference between plants grown under strong light condition, in which patterns of trichome as well as pavement cells were altered, and those grown under normal light intensity, as expected. These results indicate that the trichome pattern reflects the variation in *GL3* expression, unless the stomata pattern change, and is increased at modestly high temperatures.

**Figure 6.**
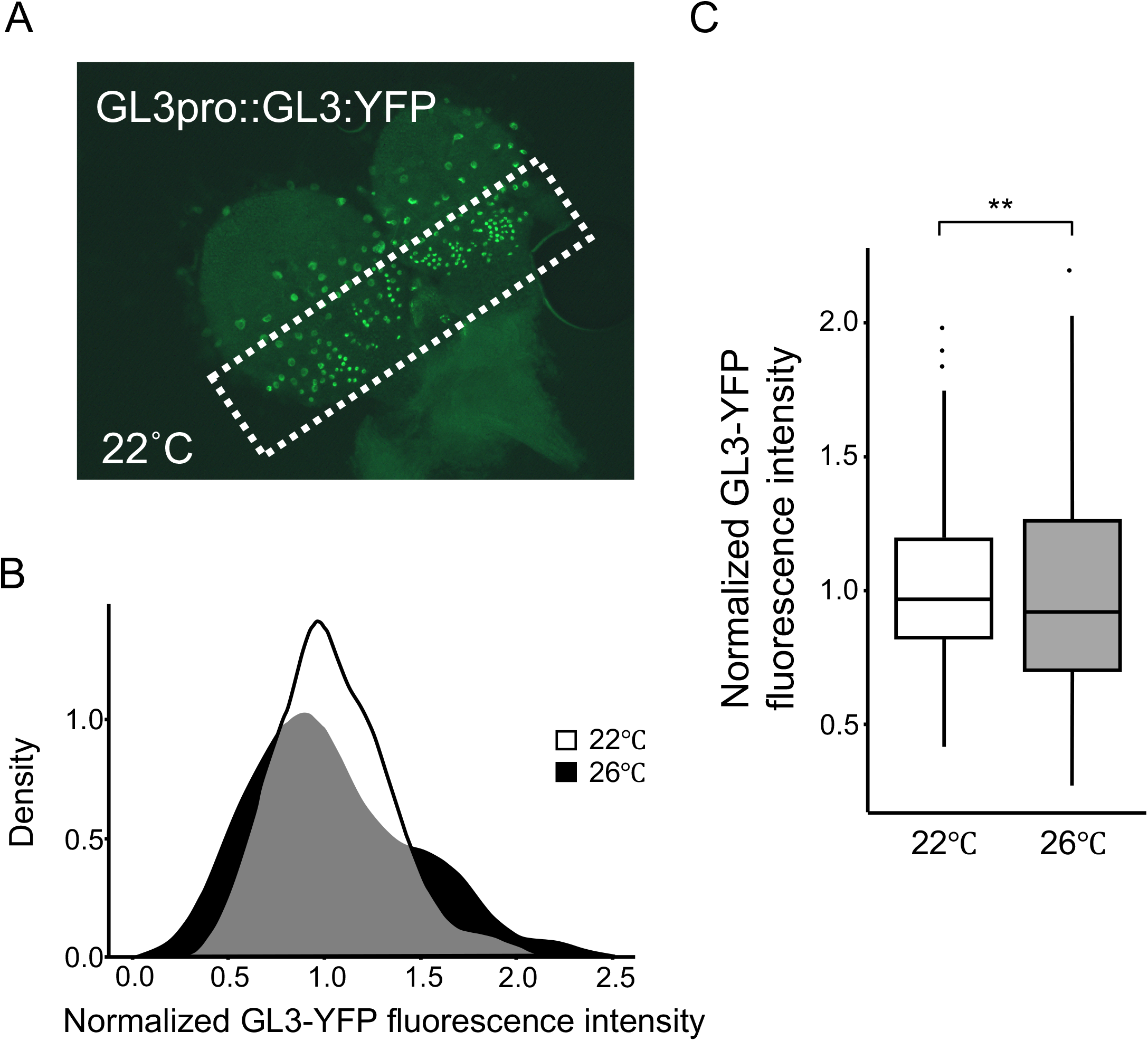
Variations in GL3 protein level. A) Representative image of GL3-YFP fluorescence signal. The GL3-YFP fluorescence intensity was measured from the area outlined by a white dotted line, which shows the initial cells in trichome development. B) Distribution of GL3-YFP fluorescence in plants grown at 22°C (white) and 26°C (black). C) Box plots of variances of intensities of normalized GL3-YFP fluorescence shown in panel B. **P < 0.01.

### 2.4 Trichome distribution patterns are variable in accessions and climate

Based on the above findings, we speculated that variations in gene expression under certain conditions could be predicted based on the regularity of the trichome pattern, and this could have an adaptive significance. We were also curious to determine which natural factors are responsible for the gene expression variation. We addressed these questions by comparing the regularity of trichome distribution patterns between of *A. thaliana* accessions adapted to the different climatic conditions [29–31]. A total of 11 accessions (Don-0, Aitba-1, Col-0, IP-Tri-0, Yo-0, Ra-0, Van-0, Ler-0, Spro-0, Pi-0, and Kin-0) were grown together in one plate under a normal or modestly high temperature (Fig. 7). Strikingly, the results indicated that the change in NDD variance varied between accessions. The accessions could be divided into two groups at 26°C: the high-variance group, comprising Don-0, Aitba-1, and Col; and the low-variance group, comprising Ler and other accessions. IP-Tri-0 was the only accession that showed no difference in NND variance distribution between at 22°C and 26°C. This led us to another question, i.e., whether gene expression variations would reflect the specific environment to which an accession is adapted.

**Figure 7.**
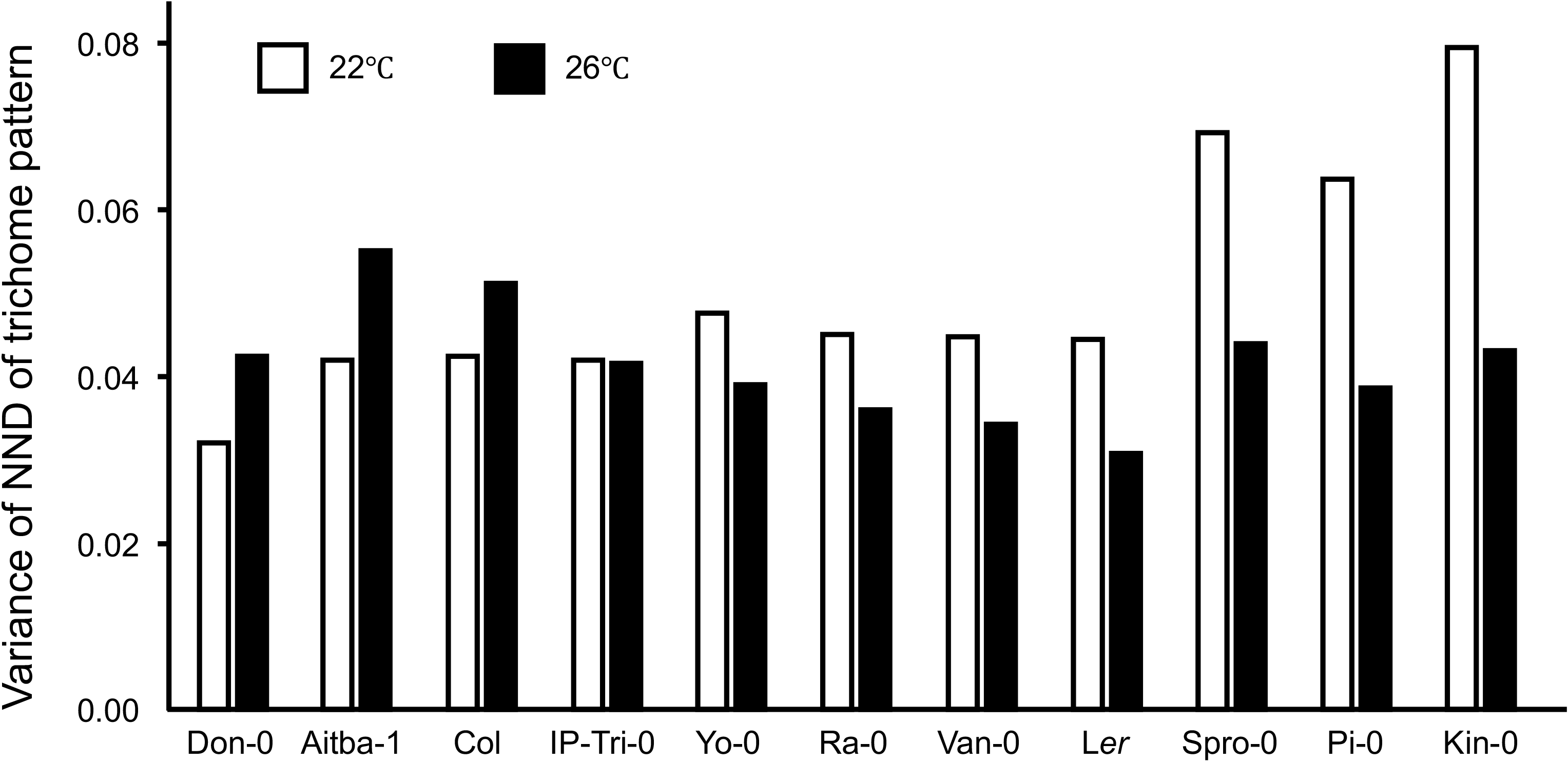
Normalized next-neighbor distance (NND) variances of trichomes of various Arabidopsis accessions grown under mild heat stress. NND variances of trichomes of Col, Ler, Van, and Kin plants grown at 22°C (white) and 26°C (black) are shown.

To evaluate the relationship between gene expression variation and environmental factors, we used the BioClim data set, which comprises 19 global land surface datasets (Table 2; Worldclim [http://www.worldclim.org/current]) [32]. Since Col and Ler have been grown under experimental conditions for more than 70 years, we excluded Col and Ler in this analysis. We calculated the ratio of variance at 26°C to that at 22°C. When gene expression variation increased under mild heat, the ratio of variances increased beyond 1.0. We analyzed the ratio of variances of 11 accessions and compared these ratios with BioClim indices (Fig. S5). We found that three indices (BIO1, BIO10, and BIO11) showed significant positive correlations with the ratio of variances of NND. Interestingly, all of these three indices represent mean temperature under certain conditions. In particular, BIO1, the index of the mean of annual temperature, was positively correlated with the variability in the trichome distribution patterns (Fig. 8). These results demonstrate a strong relationship between gene expression variation and environmental factors, especially temperature, suggesting that temperature affects gene expression variations.

**Table 2.** BioClim indices.

| BioClim Index | Description |
| --- | --- |
| BIO1 | Annual Mean Temperature |
| BIO2 | Mean Diurnal Range (Mean of monthly (max temp - min temp)) |
| BIO3 | Isothermality (BIO2/BIO7) (* 100) |
| BIO4 | Temperature Seasonality (standard deviation *100) |
| BIO5 | Max Temperature of Warmest Month |
| BIO6 | Min Temperature of Coldest Month |
| BIO7 | Temperature Annual Range (BIO5-BIO6) |
| BIO8 | Mean Temperature of Wettest Quarter |
| BIO9 | Mean Temperature of Driest Quarter |
| BIO10 | Mean Temperature of Warmest Quarter |
| BIO11 | Mean Temperature of Coldest Quarter |
| BIO12 | Annual Precipitation |
| BIO13 | Precipitation of Wettest Month |
| BIO14 | Precipitation of Driest Month |
| BIO15 | Precipitation Seasonality (Coefficient of Variation) |
| BIO16 | Precipitation of Wettest Quarter |
| BIO17 | Precipitation of Driest Quarter |
| BIO18 | Precipitation of Warmest Quarter |
| BIO19 | Precipitation of Coldest Quarter |

**Figure 8.**
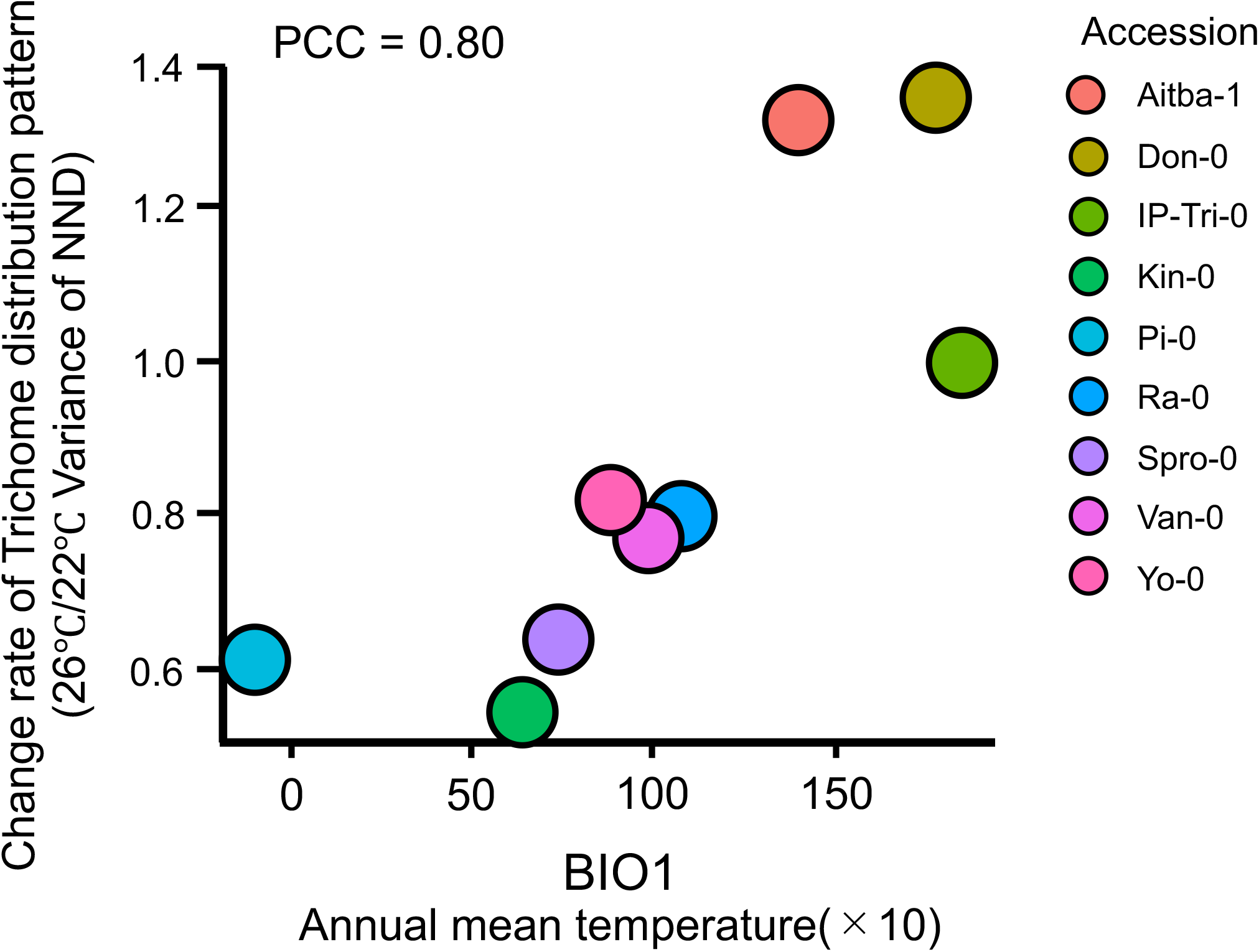
Relationship between gene expression variation and BioClim index. The ratios of NND variances of plants grown at 26°C relative to those of plants grown at 22°C are shown along the y-axis. The BIO1 index is shown along the x-axis. The values in the plots represent Pearson’s correlation coefficients. Each accession is plotted using a different color.

### 2.5. Trichome distribution patterns are altered by histone-modifying agents

It has been reported that the epigenetic status of a gene determines its expression level [33]. We therefore quantified trichome patterns by analyzing gene expression variations in leaves of plants treated with sodium butyrate (SB) or trichostatin A (TSA), both of which inhibit histone acetyltransferase activity in plants [34]. Since histone modifications are related to various biological processes, their perturbation can have pleiotropic effects. Accordingly, treatment with 5 mM SB, a standard concentration used in experiments, reduced overall plant size (Fig. S6). However, low concentrations of SB and TSA perturbed trichome but not stomata patterns (Fig. 9), indicating that modest perturbation of histone modifications has non-uniform effects on gene expression variations, which in turn increase the variability of the trichome distribution patterns.

**Figure 9.**
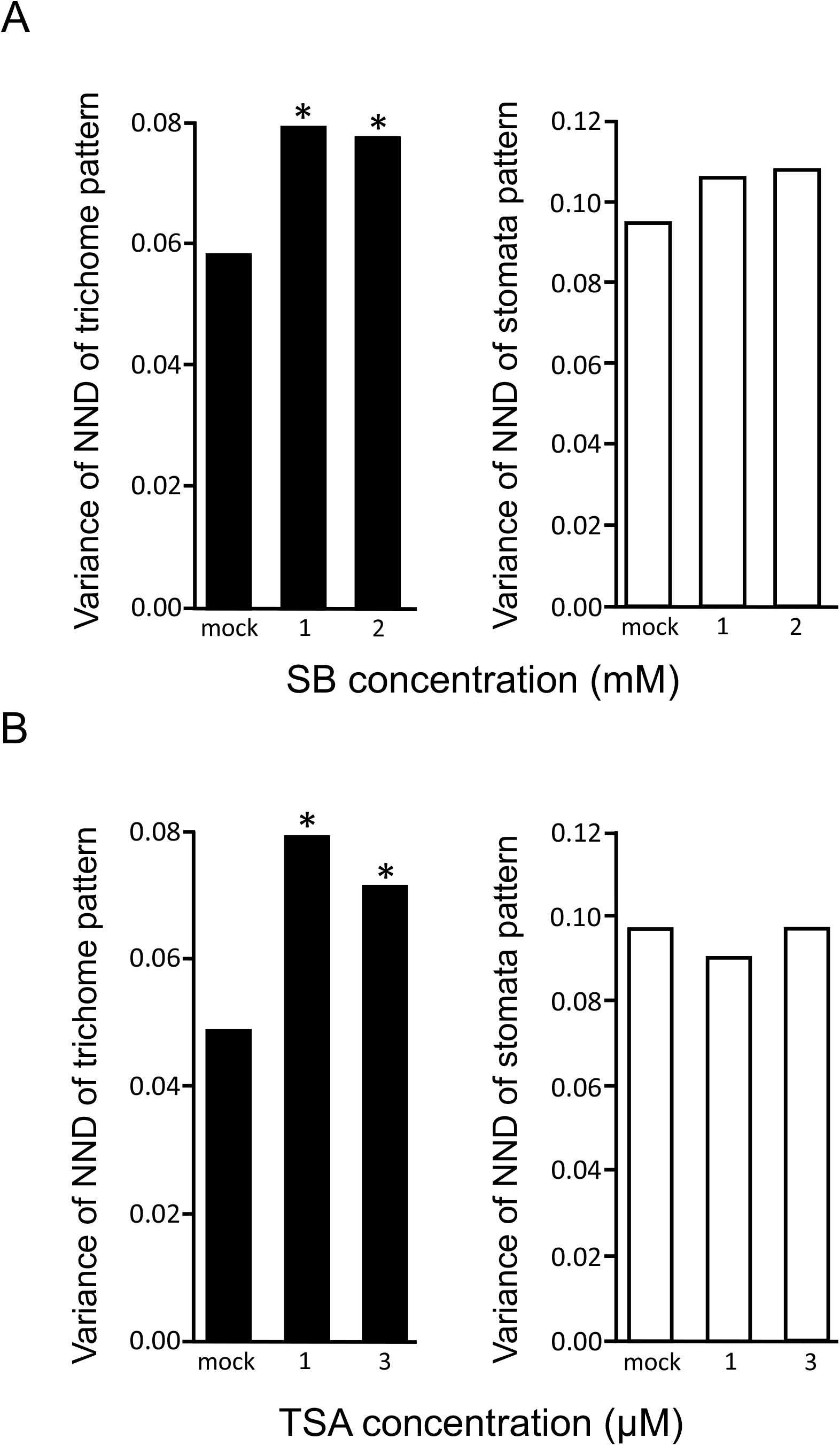
Normalized next-neighbor distance (NND) variances of trichome and stomata of Col plants treated with sodium butyrate (SB) and trichostatin A (TSA). A and B) Variances of trichomes (black) and stomata (white) of plants treated with SB (A) or TSA (B) are shown. *P < 0.05 (F test).

### 2.6. Variations in the trichome distribution patterns and H2A.Z

Since epigenetic modifications drastically alter nucleosome positions, we surveyed the nucleosome positions in the GL3 genic region using publicly available data (http://epigenomics.mcdb.ucla.edu/Nuc-Seq), which suggested that the promoter region of GL3 was relatively open (Fig S7A). To determine whether the chromatin was altered in the GL3 genic region, we performed the micrococcal nuclease (MNase) assay. No differences were observed in nucleosome positions at 22°C and 26°C (Fig. S7B). Considering our findings that the gene expression of GL3 is influenced by temperature [26], we focused on H2A.Z, which is a key histone variant involved in the thermosensing process in *Arabidopsis*. To determine whether the trichome distribution pattern is altered in the absence of a functional H2A.Z, we observed the variability in trichome distribution pattern in *arp6-1* mutant, in which eviction of H2A.Z nucleosome was perturbed [35,36]. Strikingly, Figure 10 shows that the variability in the trichome distribution pattern of the *arp6-1* mutant grown at 26°C was significantly lower than that in its background accession (Col). Moreover, the variability in the trichome distribution pattern of *arp6-1* was higher at 22°C than at 26°C.

**Figure 10.**
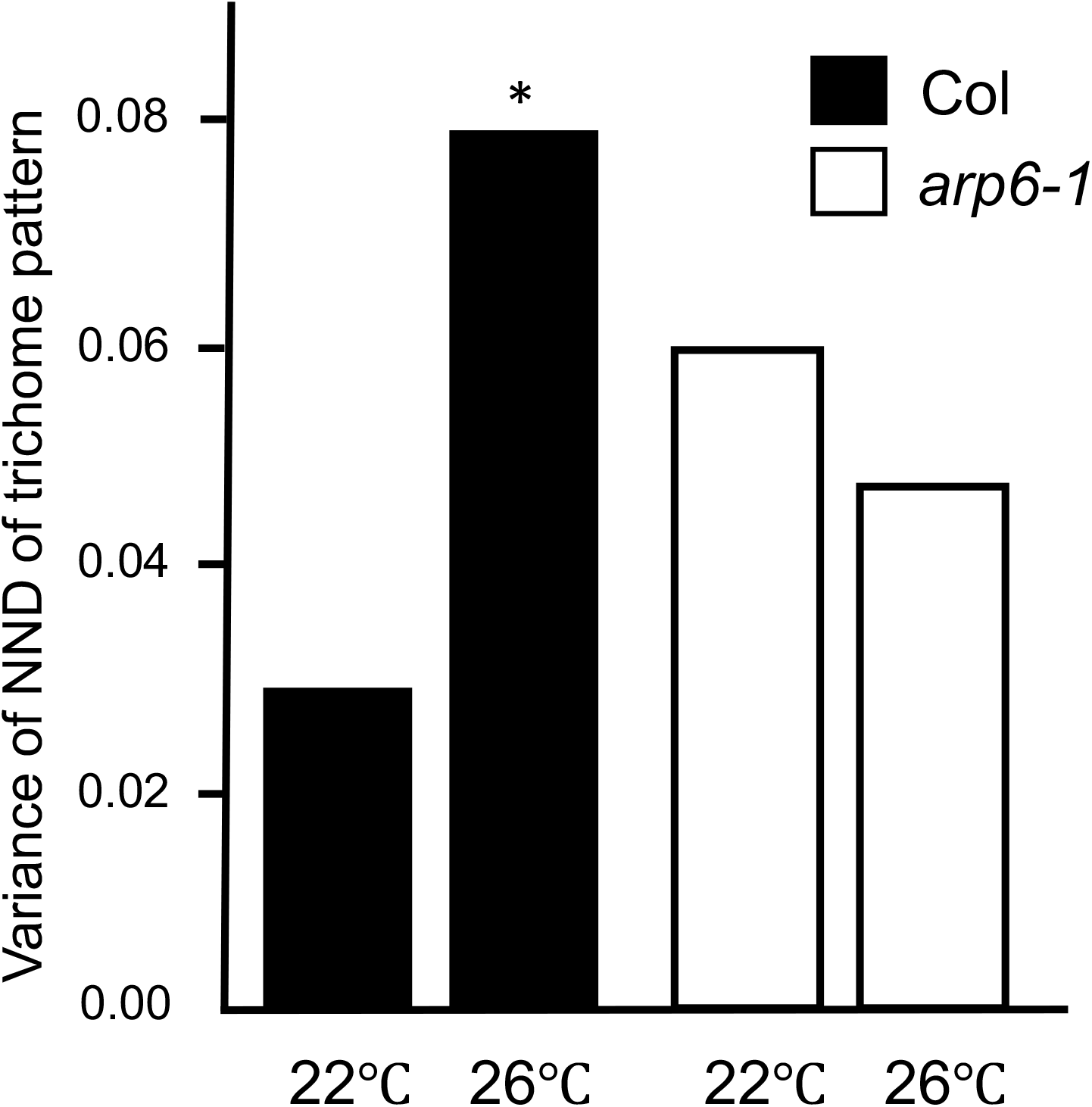
Normalized next-neighbor distance (NND) variances of trichomes in H2A.Z mutant. Variances of NND in the wild type (black) and arp6-1 (white) are shown. *P < 0.05 (F test).

## 3. Discussion

In this study, we present three major findings. First, based on the leaf trichome distribution patterns, we developed a model for measuring variations in the expression of a single gene without directly quantifying the gene expression level (e.g. by single-cell next generation sequencing). Second, the degree of gene expression variations is correlated with the annual mean temperature of the region to which the plants were adapted. Third, the H2A.Z is involved in the gene expression variation affected by mild temperature elevation.

Both GL1 and GL3 are involved in the initiation of trichome formation [14–16]; however, variations in the expression of only GL3, but not GL1, enhanced the variance of the trichome distribution pattern in our mathematical model (Fig. 3). Although it remains unclear why the variation in GL3 expression affects the trichome distribution pattern, the association of GL3 with a single MYB transcription factor (R3-MYB) may influence the variations in downstream events. Our results suggest that two parameters, β2 and γ1, decreased trichome numbers and increased the variability in the trichome distribution pattern, whereas σ2 affected only the trichome distribution pattern. Interestingly, the parameters of the other trichome factor, GL1 (σ1), and the association of GL3 with GL1 (β2), were not factors that affected the trichome distribution pattern only (Fig. S3). Our simulation also suggested that trichome numbers and patterns were independent of each other (Fig. S2). Indeed, real leaves showed no or weak correlation between trichome numbers and trichome distribution patterns (Fig. 5). Further studies are needed to investigate variations in those parameters could affect trichome numbers.

We found that changes in the trichome distribution pattern under mild heat varied between accessions; the variability in the trichome distribution pattern increased in some accessions and decreased in others under a modestly high temperature. This difference between accessions is likely attributable to the environment to which each accession is adapted. Climate is a key factor in adaptation. Trichome is thought to offer protection against herbivores, drought, and ultraviolet (UV) radiation; however, this study is the first to report that the trichome distribution pattern of a plant may contribute to its environmental fitness. Consistent with our findings of no correlation between trichome number and distribution pattern in *A. thaliana*, it has been suggested that trichomes in *A. lyrata*, are unlikely to be correlated with latitudes [37]. We speculate that to adapt, plants must be able to sense and respond to minute but significant changes in the environment. Mutations in the genome sequence are costly if the environmental change is transient; therefore, plants would have a system to quickly respond to such changes by increasing or suppressing variations in gene expression. We do not have any evidence that the trichome distribution pattern actively drives environmental adaptation in plants. The trichome pattern could be just a consequence of this response, although we cannot exclude the possibility that the trichome distribution pattern could actively contribute to plant growth in response to modest alterations in temperatures. Moreover, it is unclear whether the temperature caused genetic mutation for adaptation or a natural variation existing in each accession increased the fitness. Despite that Ler is derived from Col [38], Col and Ler showed opposite tendency with regard to the variability in trichome distribution patterns, suggesting that, at least, genetic mutations contribute the gene expression variation.

Plants employ a system comprising H2A.Z in order to sense and respond to temperature by modifying the nucleosome structure [36]. H2A.Z is a variant of histone H2A and functions to repress gene expression [39]. Ambient heat stress induces eviction of H2A.Z from the nucleosome to change gene expression. To load H2A.Z into the nucleosome requires ARP6, a subunit of the SWR1 complex. Compared with Col, *arp6-1* mutant shows increased variability of trichome distribution pattern at 22°C but similar variance at 26°C. These results suggest the involvement of H2A.Z in gene expression variation.

Variability in the trichome distribution pattern was increased by alterations in histone modifications and was correlated with the mean annual temperature. We could not generate evidence supporting altered nucleosome occupancy around the GL3 genic region (Fig. S6), possibly because of limited technical sensitivity. Other experiments, such as chromatin immunoprecipitation (ChIP) after cell sorting of trichome initial cells, may help us to reveal the cell-type specific nucleosome structure; however, it remains technically challenging. Variations in the expression of *GL3* may be more susceptible to modest environmental fluctuations than genes related to leaf development, such as those involved in pavement cell formation, and variations in gene expression could occur through epigenetic modifications including H2A.Z occupancy.

Gene expression variations are mainly classified into two categories based on the source of causal factors: intrinsic variation, which is observed intrinsically due to thermodynamics fluctuations, resulting in variable gene expression changes between cells; and extrinsic variation, which is caused by extrinsic factors, such as environmental stimuli, and is therefore observed in all cells simultaneously. In our experiments, we could not determine whether intrinsic or extrinsic variation caused the variability in the trichome distribution pattern. Variation in the nucleosome structures due to epigenetic modifications is supposedly a major contributor to intrinsic variation [2]. Our results of the chemical perturbation of histone modification revealed changes in trichome distribution patterns, which could be an evidence of intrinsic variation in the case of trichome pattern. On the other hand, altered trichome distribution patterns due to a modest heat treatment may be an example of extrinsic variation. Therefore, our approach consists of both intrinsic and extrinsic variations.

In conclusion, our finding of a correlation between the trichome distribution pattern and temperature suggests that natural variation in gene expression is associated with adaptation to the environment. Further investigation would help to understand the biological relevance of the trichome distribution pattern, as well as the mechanism underlying gene expression variation to achieve environmental adaptation.

## 4. Materials and Methods

### 4.1. Plant materials and growth conditions

*Arabidopsis thaliana* accessions Aitba-1, Col-0, Ler-0, Don-0, IP-Tri-0, Kin-0, Pi-0 Ra-0, Spro-0, Van-0, and Yo-0, obtained from the Arabidopsis Biological Resource Center (ABRC; Ohio State University, Columbus, OH, USA), were used in this study. We show data for Col and Ler, which originated from Col-0 and Ler-0, respectively. Seeds of *rol1-1* and *rol1-2* mutants were obtained from ABRC. Seeds of the *ap6-1* mutant were kindly provided by P. Wigge (John Innes Centre, UK) and K. Sugimoto (RIKEN, Japan).

Seeds were sterilized for 6 min in a solution of 50% commercial bleach (Kao, Singapore) containing 6% sodium hypochlorite and then were washed three times with distilled water. The sterilized seeds were laid out on a sterile filter paper or in 50% Murashige and Skoog (MS) medium (Wako Pure Chemical Industries, Osaka, Japan) supplemented with 1% sucrose (Wako Pure Chemical Industries) and 6% gellan gum (pH 5.9; Wako Pure Chemical Industries), and sterilized 0.1% agarose solution was added to the seeds. Plates containing the seeds were incubated at 4°C in the dark for 3 days. Subsequently, the plates were transferred to a growth chamber and incubated for 21 days at 22°C under constant light (1,000 lm/m^2^).

To conduct environmental stress treatments, plants were grown for 21 days at 22°C, 26°C, or 30°C under constant light (1,000 lm/m^2^). To perform heat stress treatments, plates were incubated at 26°C or 30°C under constant light (1,000 lm/m^2^). To perform strong light stress treatments, plants were incubated at 22°C under constant light (1,000 lm/m^2^) and strong light (3,000 lm/m^2^) after vernalization. To conduct the histone deacetylase inhibitor treatment, the filter paper on which plants were grown was transferred to a solution of trichostatin A (T8552; Sigma-Aldrich, St. Louis, MO, USA) or sodium butyrate (B5887; Sigma-Aldrich) at various concentrations followed by soaking on 50% MS medium for 1 h. The filter paper was then transferred to the original MS medium plate and cultured under the original conditions for 14 days.

### 4.2. CV of gene expression variations

Publicly available single-cell RNA sequencing data were used [20]. Since the available data consist of FPKM values that classified into cell types, coefficient variance (CV) of FPKM of each gene within a cell type was calculated. Since a low expression tends high variation, the FPKM values higher than average were used. After calculation of CV of each gene, a mean CV was calculated from whole CVs.

### 4.3. Quantification of the trichome distribution pattern

*Arabidopsis* leaves were examined by PLM as previously described [21], with minor modifications. After 21 days of growth, the third or fourth leaves were harvested, cleared, and incubated overnight in 95% ethanol and then for 1 h in 80% lactic acid. Images of cleared leaves were acquired by PLM, and the positions of trichomes were automatically marked using the ImageJ software (National Institutes of Health, Bethesda, MD, USA) with a custom script. Distances between trichomes were calculated using the R software (31), and NNDs were calculated according to the following equation:

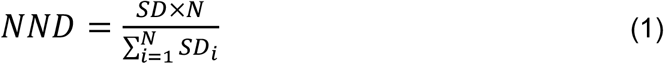

where *SD_i_* is the *i^th^* shortest path between two trichomes, and *N* is the number of trichomes. The variance of NND was calculated, and the F test was used to validate the significance of differences between variance values. At least 125 trichomes derived from five or more plants were examined. To analyze the stomata pattern, leaves were stained with PI and observed by fluorescence microscopy. The NNDs of stomata were calculated using the equation (1) but with stomata positions, not trichome positions. At least 306 stomata derived from five or more plants were analyzed.

### 4.4. Estimation of the range of biologically appropriate stochastic variations

To determine the range of biologically appropriate variations, the stochastic parameters, *K_v_*, were calculated using the following equation:

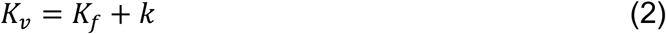

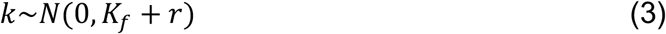

where *K_f_* is a fixed value from Table 1, and *r* ranges from 0% to 50% of the value of *K_f_*. The symbol *K_v_* was assigned to all parameters, as long as trichome patterns appeared in the simulation.

### 4.5. Mathematical modeling and simulations

All simulations in this study were performed using Matlab (9.0.0.341360 [R2016a]; MathWorks, Natick, MA, USA). The mathematical model and scripts were based on a previous study [13], with some modifications. Based on the interaction diagram shown in Figure 1A, the following equations were used:

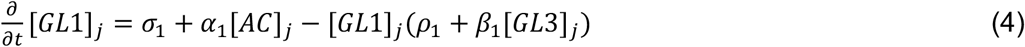

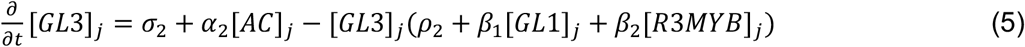

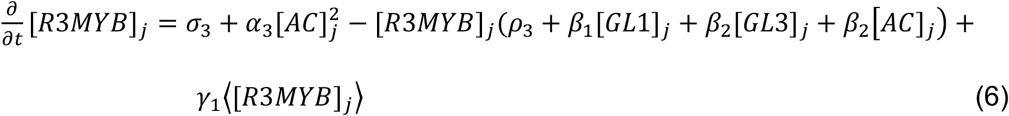

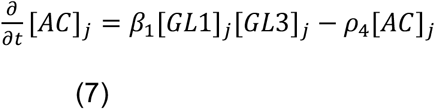

The values of parameters used are shown in Table 1. The steady state was determined with an additional 1% stochastic variation in parameters per cell as the initial condition for the simulation of cell grids. Simulations were performed 500 times using 10,000 grids, unless specified otherwise.

### 4.6. Comparison of climate data

To compare the regularity of trichome distribution patterns with respect to climate data, 19 BioClim indices (BIO1–19) [32] were used (Table 2). To obtain correlation coefficients between these indices and determine the regularity of the trichome distribution pattern, the ratio of variances between plants grown at 22°C and 26°C were calculated, yielding a dimensionless value, which was plotted against each BioClim index. Then, the Pearson’s correlation coefficient was calculated.

### 4.7. MNase assay

The micrococcal nuclease (MNase) assay was performed according to [40]. 0.5 g of 3-weeks seedlings were collected, frozen with liquid nitrogen, and crushed in a tube with beads and a crusher. 10 ml nuclear extraction buffer A (0.25 M sucrose, 60 mM KCl, 15 mM MgCl_2_, 1 mM CaCl_2_, 15 mM PIPES (pH 6.8), 0.8% Triton X-100, and 1 mM phenylmethylsulfonyl fluoride (PMSF)) was added to the crushed tissue in a 50 ml Falcon tube. The crushed tissue sample was mixed with a vortex and filtered through a funnel with a single layer of Miracloth (MilliporeSigma, Burlington, Massachusetts). The sample was filtered twice in total. After centrifugation at 10,000 g for 20 minutes at 4°C, the supernatant was removed, and 500 μl of nuclear extraction buffer B (0.25 M sucrose, 10 mM Tris-HCl (pH 8.0), 10 mM MgCl_2_, 1% Triton X-100, 5 mM 2-mercaptoethanol, and 1 mM PMSF) was added and resuspended. An equal amount of nuclear extraction buffer C (1.7 M sucrose, 10 mM Tris-HCl (pH 8.0), 10 mM MgCl_2_, 0.5% Triton X-100, 5 mM 2-mercaptoethanol, and 1 mM PMSF) was prepared in a new 1.5 ml tube and the resuspended sample was gently added. After centrifugation at 12,000 g for 1 hour at 4°C, all the supernatant was removed and resuspended in 250 μl MNase buffer (0.3 M sucrose, 20 mM Tris-HCl (pH 7.5), and 3 mM CaCl_2_). The DNA concentration at this time was measured. After adjusting the DNA concentration with MNase buffer within 300-600 ng/μl, 10 μl MNase (2,000,000 gels units/ml, NEB, Ipswich, Massachusetts) with appropriate units was added to 30 μl sample and incubated at 37°C for 15 minutes. After incubation, 40 μl of 2x stop buffer (50 mM EDTA and 1% SDS), 10x proteinase K buffer (100 mM Tris-HCl (pH 7.8), 50 mM EDTA, and 5% SDS) and 1 μl of proteinase K (800 units/ml) were added and incubated at 45°C for 1 hour. The DNA was purified by the phenol/chloroform extraction, followed by the ethanol precipitation. After the fragmented DNA concentration were confirmed by agarose gel electrophoresis, real-time PCR was performed. *At4g07700*, a gypsy-like transposable element, was used as a reference in real-time PCR [41]. The oligo primers used are shown in Table 3.

**Table 3.**
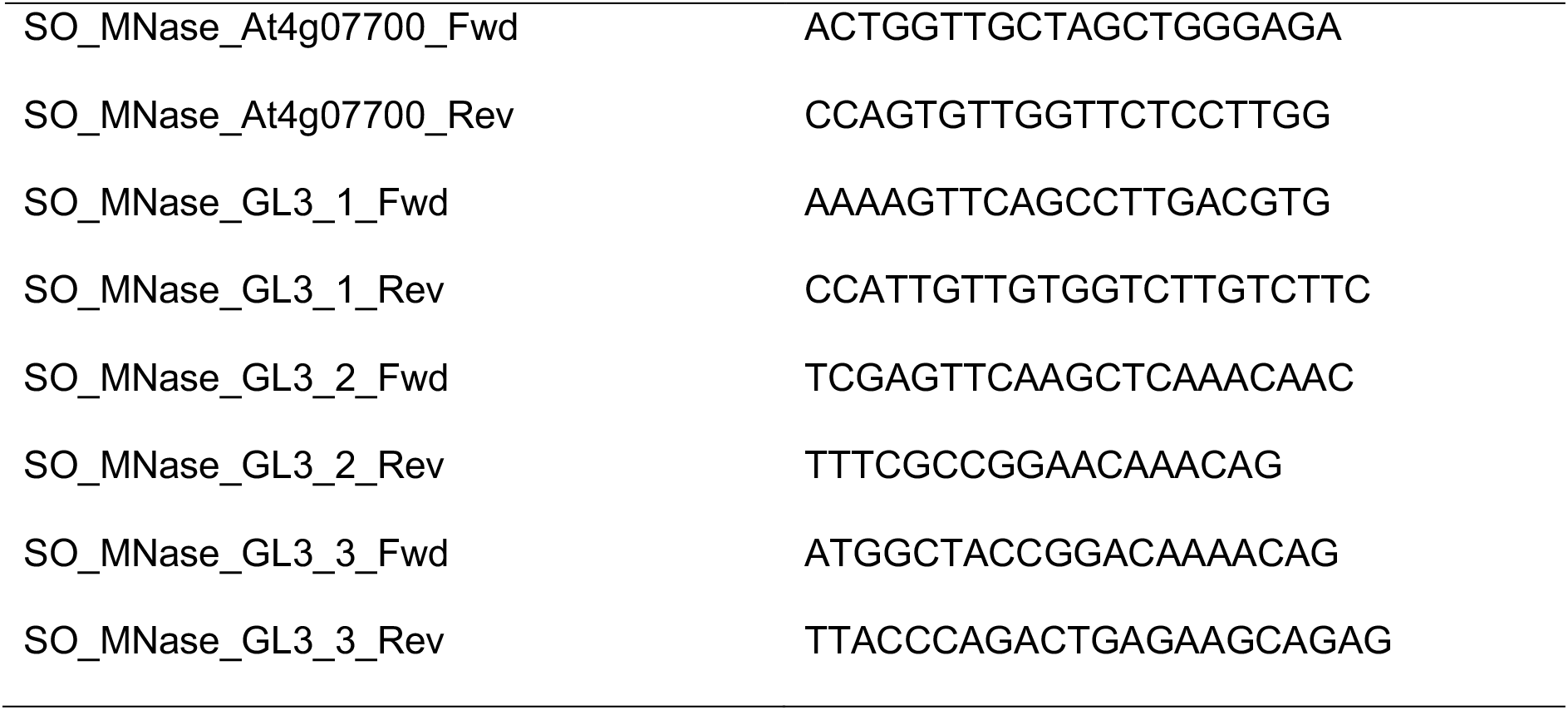
Oligo primers used for MNase assay.

## Supplementary Materials

**Figure S1:** Coefficient of variation (CV) of gene expression variations; **Figure S2**: Relationship between the number of trichomes and regularity of the trichome distribution pattern; **Figure S3:** Variances of the normalized next-neighbor distance (NND) and number of trichomes in mathematical simulations; **Figure S4:** NND variances of the trichomes and stomata of *Arabidopsis* plants grown under different light intensities; **Figure S5:** Correlation between gene expression variation and BioClim indices; **Figure S6:** Size of seedlings grown in the presence of various concentrations of sodium butyrate, a histone deacetylase inhibitor; **Figure S7**: Micrococcal nuclease (MNase) assay.

## Author Contributions

Conceptualization, S.O. and K.M.; methodology, S.O., T.U. and K.M.; validation, S.O., K.N. and T.Y.; formal analysis, S.O., K.N., and Y.T.; investigation, S.O. and Y.T; data curation, S.O. T.U. and K.M.; writing—original draft preparation, K.M.; writing—review and editing, T.U. and K.M..; supervision, K.M.; project administration, K.M.; funding acquisition, K.M. All authors have read and agreed to the published version of the manuscript.

## Acknowledgments

We thank P. Wigge (John Innes Centre, UK) and K. Sugimoto (RIKEN, Japan) for sharing *arp6-1* mutant seeds. We also thank V. Teva for critically reading the manuscript and providing useful suggestions, and H. Takemura for helpful discussions and suggestions regarding the simulation scripts. We thank N. Takayanagi for providing invaluable assistance in conducting the experiments.

