## Supplemental figures for "Leaf trichome distribution pattern in *Arabidopsis* reveals gene expression variation associated with environmental adaptation": Figures_S_ver20.pptx

### Slide 1
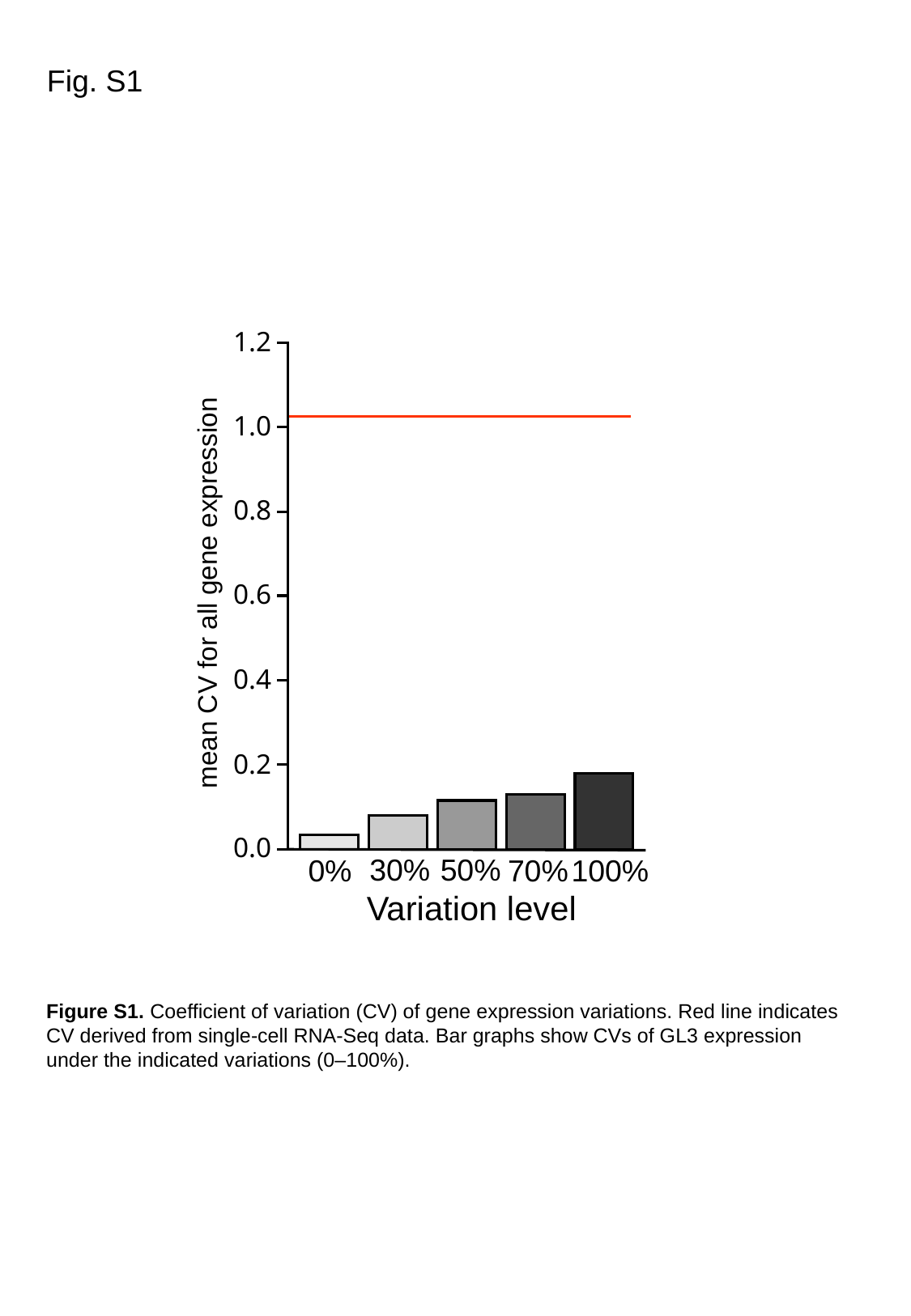

Fig. S1
1.2
1.0
0.8
mean CV for all gene expression
0.6
0.4
0.2
0.0
50%
30%
0%
70%
100%
Variation level
Figure S1. Coefficient of variation (CV) of gene expression variations. Red line indicates CV derived from single-cell RNA-Seq data. Bar graphs show CVs of GL3 expression under the indicated variations (0–100%).

### Slide 2
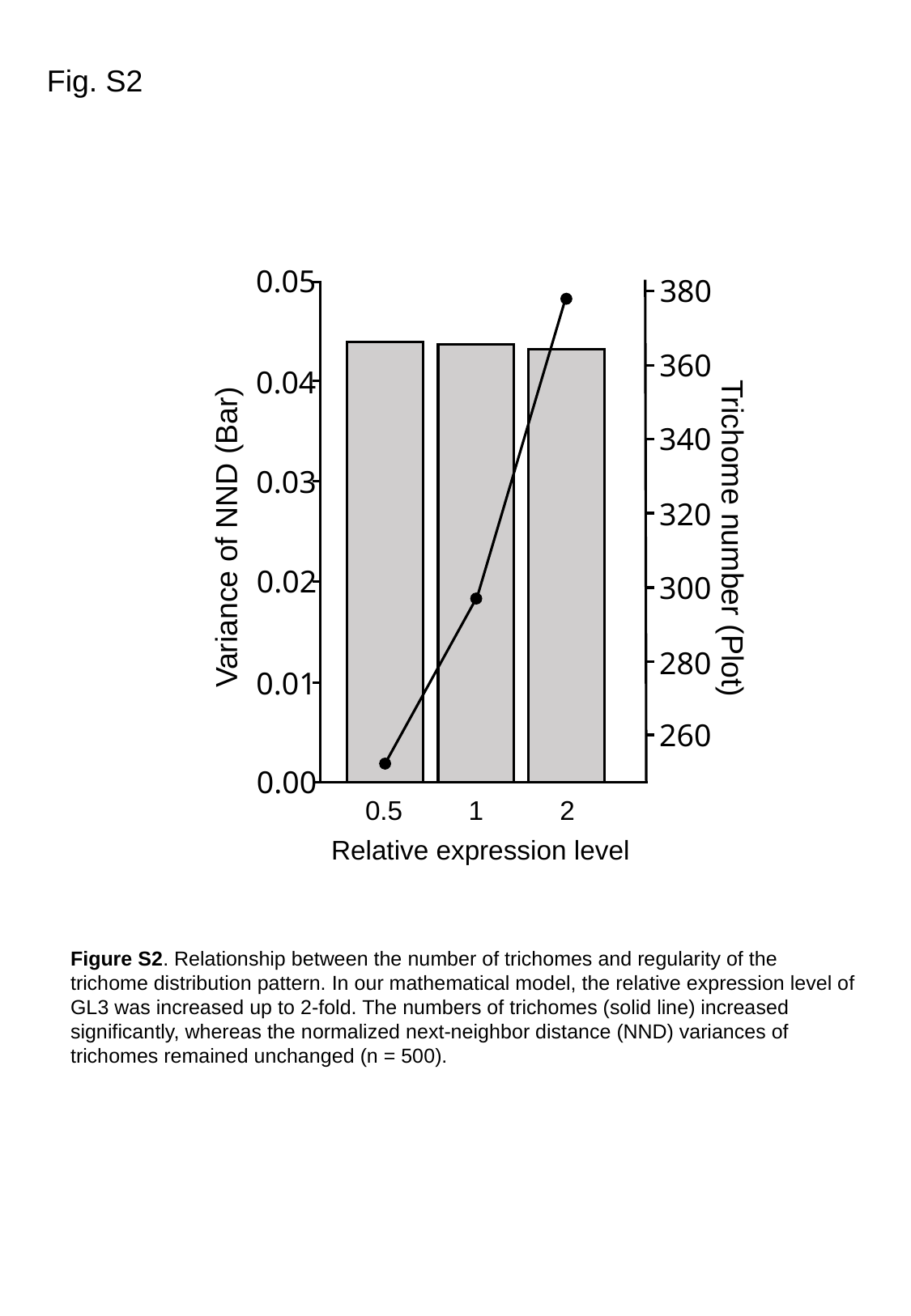

Fig. S2
0.05
380
360
0.04
340
0.03
320
Variance of NND (Bar)
Trichome number (Plot)
0.02
300
280
0.01
260
0.00
0.5
1
2
Relative expression level
Figure S2. Relationship between the number of trichomes and regularity of the trichome distribution pattern. In our mathematical model, the relative expression level of GL3 was increased up to 2-fold. The numbers of trichomes (solid line) increased significantly, whereas the normalized next-neighbor distance (NND) variances of trichomes remained unchanged (n = 500).

### Slide 3
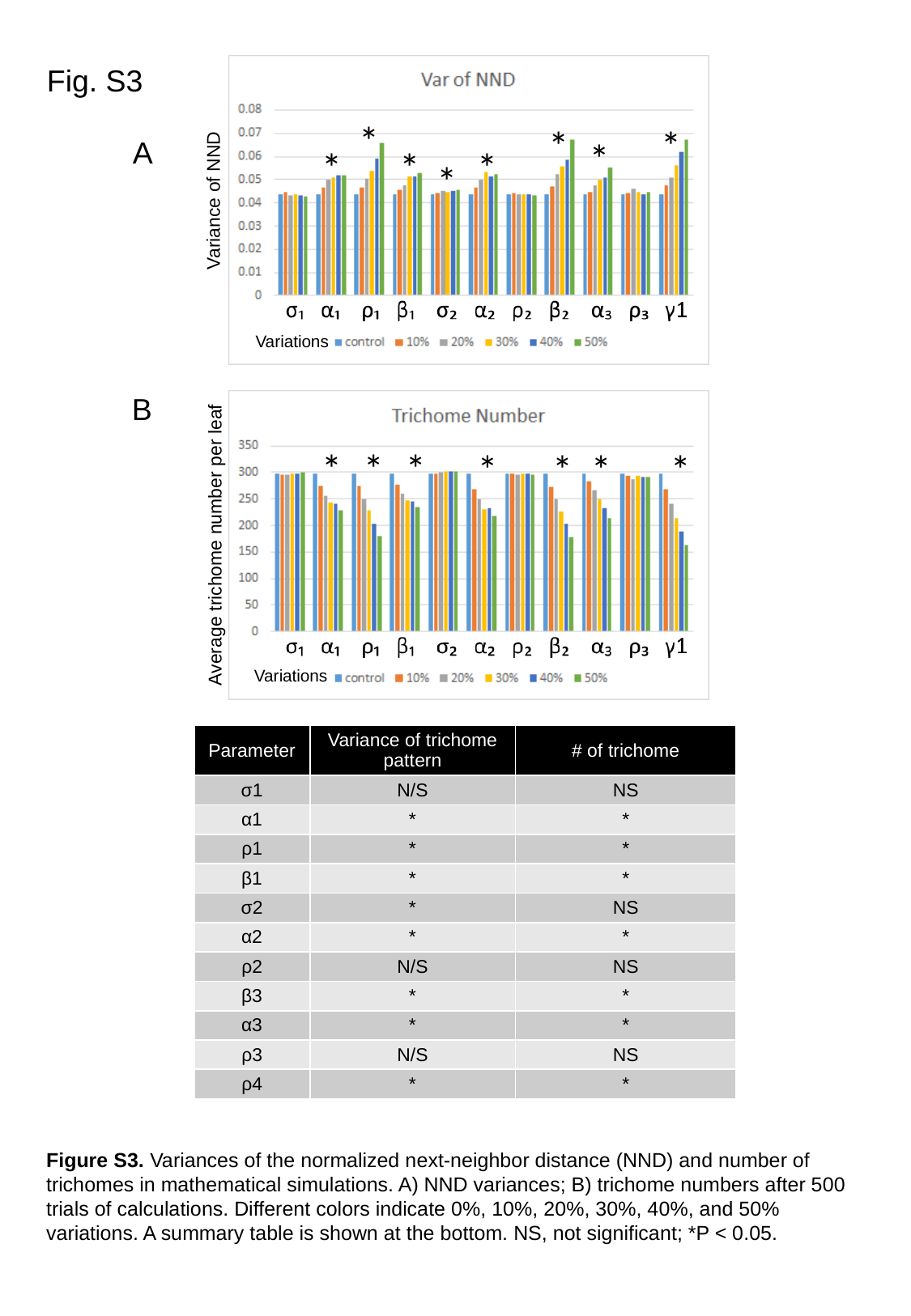

Fig. S3
*
*
*
*
*
*
*
*
Variance of NND
Variations
*
*
*
*
*
*
*
Average trichome number per leaf
Variations
A
B
| Parameter | Variance of trichome pattern | # of trichome |
| --- | --- | --- |
| σ1 | N/S | NS |
| α1 | \* | \* |
| ρ1 | \* | \* |
| β1 | \* | \* |
| σ2 | \* | NS |
| α2 | \* | \* |
| ρ2 | N/S | NS |
| β3 | \* | \* |
| α3 | \* | \* |
| ρ3 | N/S | NS |
| ρ4 | \* | \* |
Figure S3. Variances of the normalized next-neighbor distance (NND) and number of trichomes in mathematical simulations. A) NND variances; B) trichome numbers after 500 trials of calculations. Different colors indicate 0%, 10%, 20%, 30%, 40%, and 50% variations. A summary table is shown at the bottom. NS, not significant; *P < 0.05.

### Slide 4
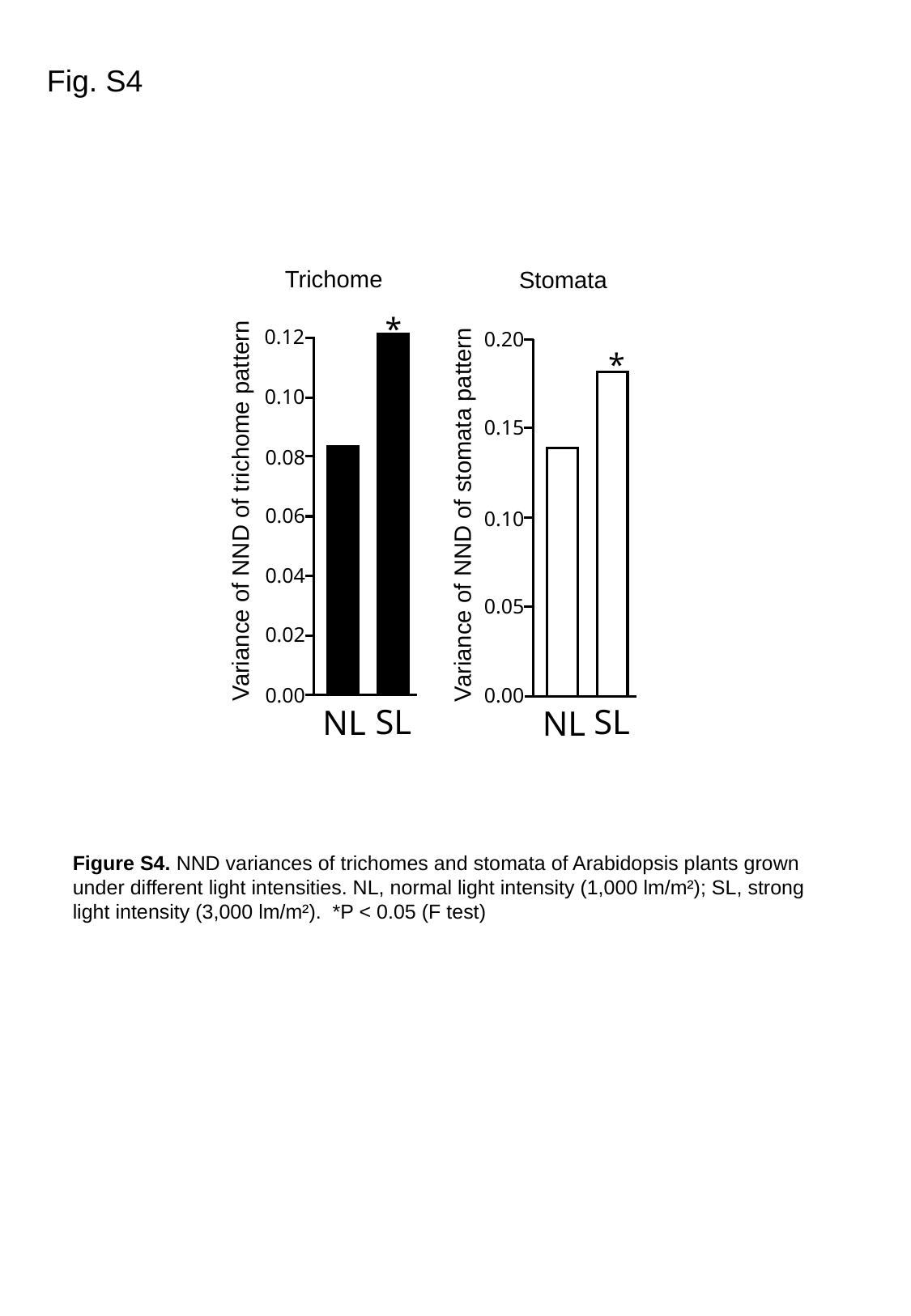

Fig. S4
Trichome
Stomata
*
0.12
0.10
0.08
Variance of NND of trichome pattern
0.06
0.04
0.02
0.00
SL
NL
0.20
*
0.15
Variance of NND of stomata pattern
0.10
0.05
0.00
SL
NL
Figure S4. NND variances of trichomes and stomata of Arabidopsis plants grown under different light intensities. NL, normal light intensity (1,000 lm/m²); SL, strong light intensity (3,000 lm/m²). *P < 0.05 (F test)

### Slide 5
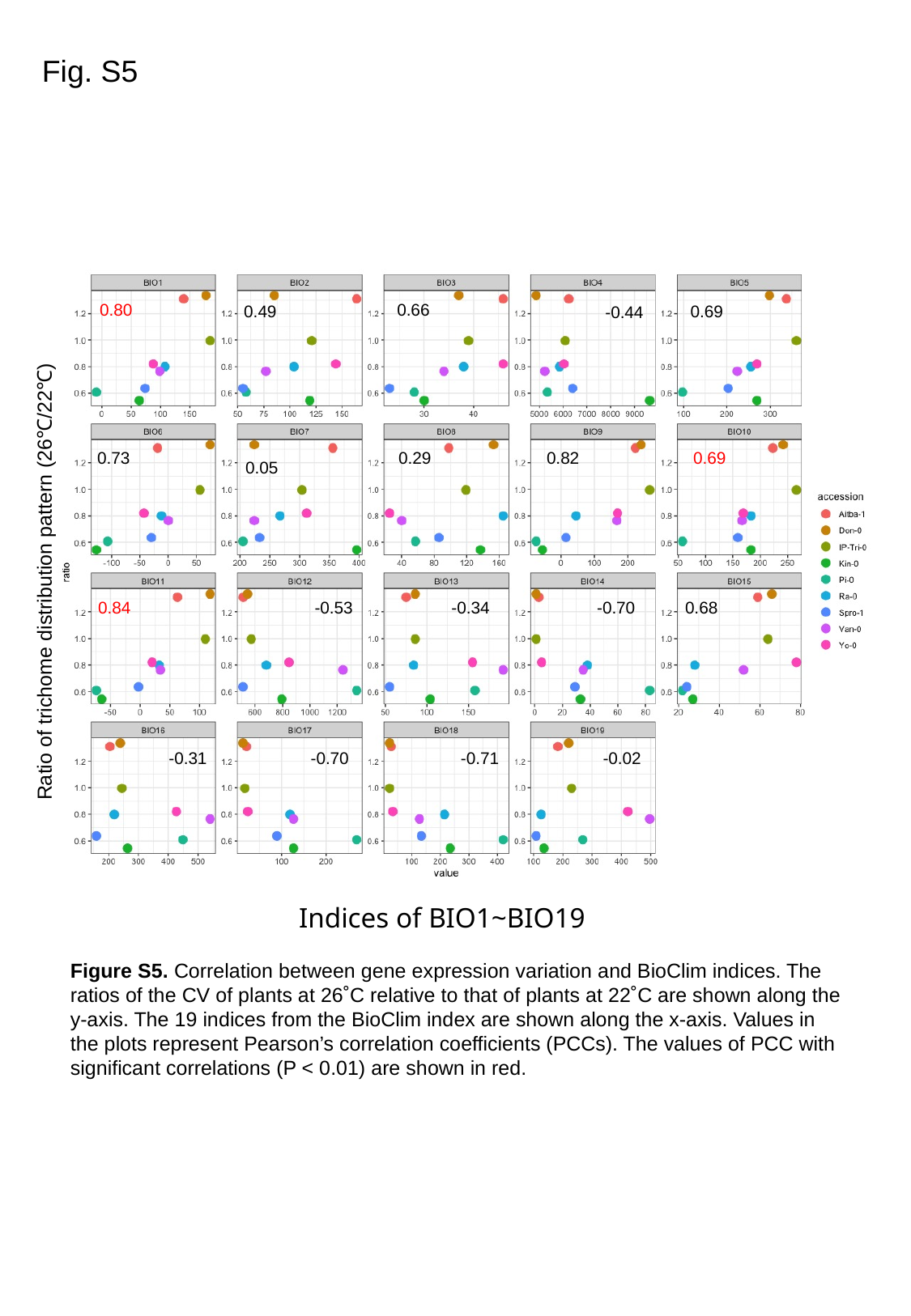

Fig. S5
0.80
0.66
0.49
0.69
-0.44
0.73
0.29
0.82
0.69
0.05
Ratio of trichome distribution pattern (26℃/22℃)
0.84
-0.53
-0.34
-0.70
0.68
-0.31
-0.70
-0.71
-0.02
Indices of BIO1~BIO19
Figure S5. Correlation between gene expression variation and BioClim indices. The ratios of the CV of plants at 26˚C relative to that of plants at 22˚C are shown along the y-axis. The 19 indices from the BioClim index are shown along the x-axis. Values in the plots represent Pearson’s correlation coefficients (PCCs). The values of PCC with significant correlations (P < 0.01) are shown in red.

### Slide 6
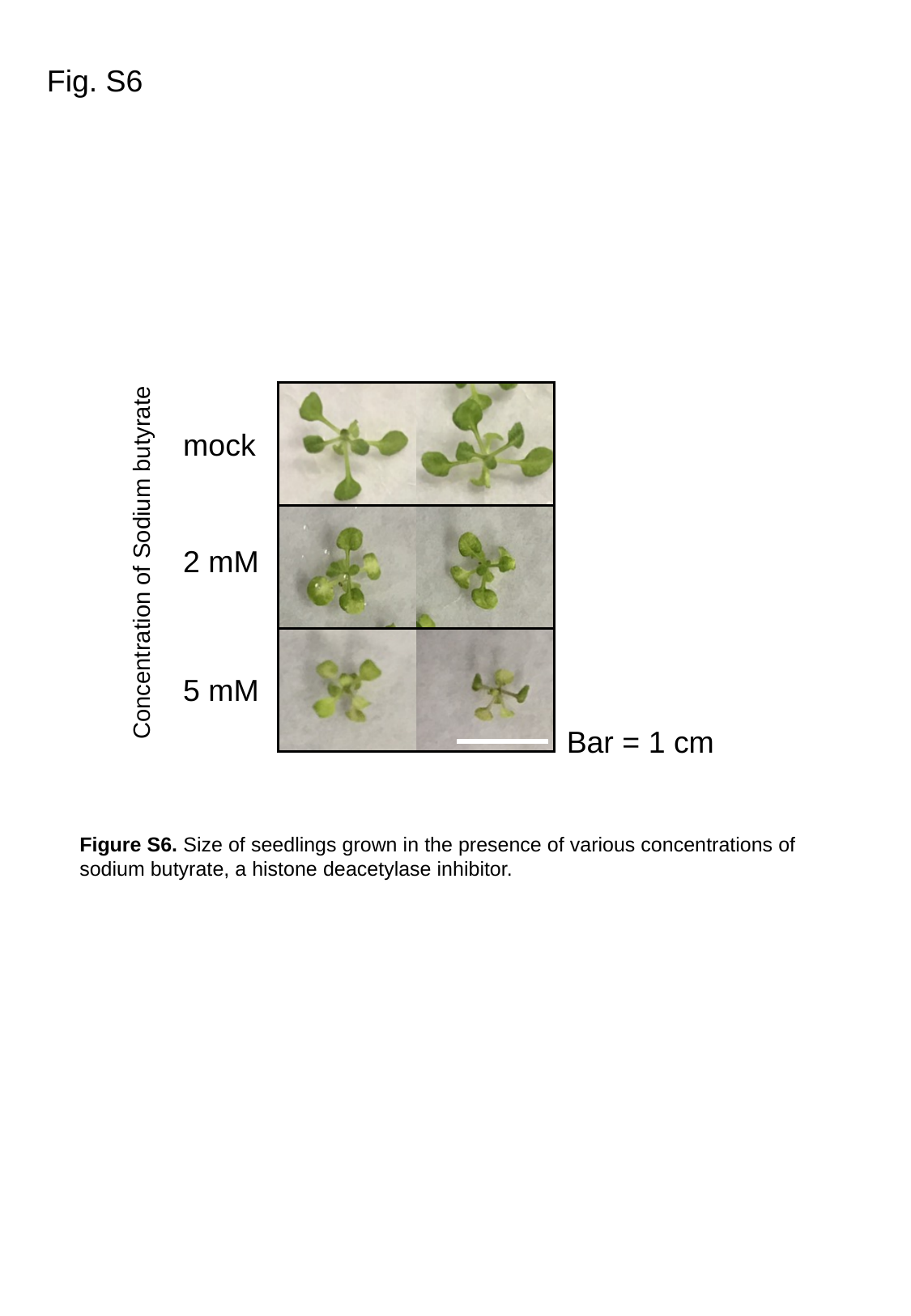

Fig. S6
mock
2 mM
Concentration of Sodium butyrate
5 mM
Bar = 1 cm
Figure S6. Size of seedlings grown in the presence of various concentrations of sodium butyrate, a histone deacetylase inhibitor.

### Slide 7
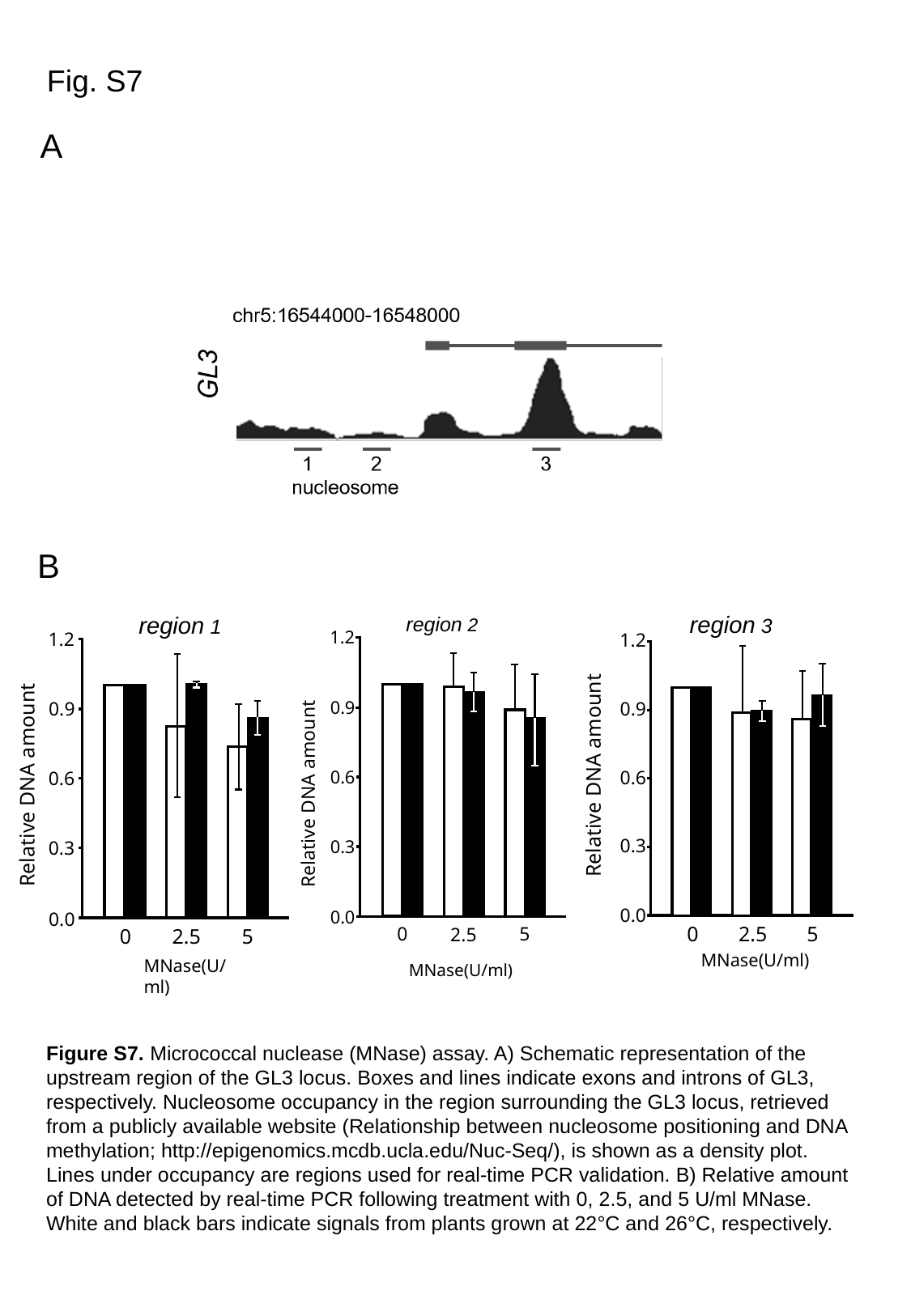

Fig. S7
A
B
region 3
1.2
0.9
0.6
0.3
0.0
0
5
2.5
MNase(U/ml)
Relative DNA amount
region 1
1.2
0.9
0.6
0.3
0.0
0
5
2.5
MNase(U/ml)
Relative DNA amount
region 2
1.2
0.9
0.6
0.3
0.0
0
5
2.5
MNase(U/ml)
Relative DNA amount
Figure S7. Micrococcal nuclease (MNase) assay. A) Schematic representation of the upstream region of the GL3 locus. Boxes and lines indicate exons and introns of GL3, respectively. Nucleosome occupancy in the region surrounding the GL3 locus, retrieved from a publicly available website (Relationship between nucleosome positioning and DNA methylation; http://epigenomics.mcdb.ucla.edu/Nuc-Seq/), is shown as a density plot. Lines under occupancy are regions used for real-time PCR validation. B) Relative amount of DNA detected by real-time PCR following treatment with 0, 2.5, and 5 U/ml MNase. White and black bars indicate signals from plants grown at 22°C and 26°C, respectively.
